# EWS-FLI-1 creates a cell surface microenvironment conducive to IGF signaling by inducing pappalysin-1

**DOI:** 10.1101/195693

**Authors:** Panneerselvam Jayabal, Peter J. Houghton, Yuzuru Shiio

**Affiliations:** Greehey Children’s Cancer Research Institute, The University of Texas Health Science Center, San Antonio, Texas 78229-3900, USA; Cancer Therapy and Research Center, The University of Texas Health Science Center, San Antonio, Texas 78229-3900, USA; Department of Molecular Medicine, The University of Texas Health Science Center, San Antonio, Texas 78229-3900, USA; Department of Biochemistry, The University of Texas Health Science Center, San Antonio, Texas 78229-3900, USA

**Keywords:** Ewing sarcoma, EWS-FLI-1, IGFBP, IGF signaling, pappalysin-1

## Abstract

Ewing sarcoma is an aggressive cancer of bone and soft tissue in children with poor prognosis. It is characterized by the chromosomal translocation between EWS and an Ets family transcription factor, most commonly FLI-1. EWS-FLI-1 fusion accounts for 85% of Ewing sarcoma cases. EWS-FLI-1 regulates the expression of a number of genes important for sarcomagenesis, can transform NIH3T3 and C3H10T1/2 cells, and is necessary for proliferation and tumorigenicity of Ewing sarcoma cells, suggesting that EWS-FLI-1 is the causative oncoprotein.

Here we report that EWS-FLI-1 induces the expression of pappalysin-1 (PAPPA), a cell surface protease that degrades IGF binding proteins (IGFBPs) and increases the bioavailability of IGF. EWS-FLI-1 binds to the pappalysin-1 gene promoter and stimulates the expression of pappalysin-1, leading to degradation of IGFBPs and enhanced IGF signaling. Silencing of pappalysin-1 strongly inhibited anchorage-dependent and anchorage-independent growth as well as xenograft tumorigenicity of Ewing sarcoma cells. These results suggest that EWS-FLI-1 creates a cell surface microenvironment conducive to IGF signaling by inducing pappalysin-1, which emerged as a novel target to inhibit IGF signaling in Ewing sarcoma.

## Introduction

Ewing sarcoma is an aggressive cancer of bone and soft tissue in children with poor long-term outcome. Ewing sarcoma is characterized by the reciprocal chromosomal translocation generating a fusion oncogene between EWS and an Ets family transcription factor, most commonly FLI-1 [1-5]. EWS-FLI-1 translocation accounts for 85% of Ewing sarcoma cases. The EWS-FLI-1 gene product regulates the expression of a number of genes important for cancer progression [6], can transform mouse cells such as NIH3T3 [7] and C3H10T1/2 [8], and is necessary for proliferation and tumorigenicity of Ewing sarcoma cells [1-5]. These findings suggest that EWS-FLI-1 plays a central role in Ewing sarcomagenesis although the role for the reciprocal fusion, FLI-1-EWS, was also suggested recently [9].

IGF signaling plays an important role in Ewing sarcomagenesis. IGF-1 and IGF-1R expression are elevated in Ewing sarcoma cell lines and tumors [10, 11]. Blockage of IGF-1R by a monoclonal antibody inhibits the xenograft tumorigenicity of Ewing sarcoma cells [12]. IGF-1R expression is necessary for transformation of mouse fibroblasts by EWS-FLI-1 [13]. Expression of EWS-FLI-1 in mesenchymal stem cells, the putative cells of origin for Ewing sarcoma [1-5], results in the induction of IGF-1 and IGF-1 dependence for cell growth [14, 15]. EWS-FLI-1 directly represses the expression of IGFBP3 [16], which functions as an inhibitor of IGF signaling by binding and sequestering IGFs.

Using secretome proteomics, we found that EWS-FLI-1 induces the expression of pappalysin-1, a cell surface protease that cleaves IGFBP2, IGFBP4, and IGFBP5 [17, 18]. Like IGFBP3, IGFBP2, IGFBP4, and IGFBP5 function as inhibitors of IGF signaling. We demonstrate that EWS-FLI-1 binds to the gene promoter of pappalysin-1 and activates its expression. Silencing of pappalysin-1 resulted in accumulation of IGFBPs and reduced bioactive IGF-1 in Ewing sarcoma cell secretome, leading to suppression of IGF signaling. Pappalysin-1 silencing strongly inhibited both anchorage-dependent and anchorage-independent growth as well as xenograft tumorigenicity of Ewing sarcoma cells. These results suggest the critical role played by the EWS-FLI-1 – pappalysin-1 axis in creating a cell surface microenvironment that facilitates IGF signaling in Ewing sarcoma.

## Results & Discussion

### EWS-FLI-1 activates pappalysin-1 expression in Ewing sarcoma

To dissect the impact of EWS-FLI-1 on Ewing sarcoma cell secretome, we silenced EWS-FLI-1 using lentiviruses expressing an shRNA against FLI-1 C-terminal region in A673 Ewing sarcoma cells (Fig. 1A; luciferase shRNA as control) and analyzed the proteins secreted in the conditioned medium by GeLC-MS/MS and spectral counting. (The details of the proteomic analysis will be published elsewhere.) From this, we found dramatically reduced pappalysin-1 protein levels in A673 cell secretome upon EWS-FLI-1 silencing (Fig. 1B). EWS-FLI-1 silencing also reduced pappalysin-1 transcript levels in A673 cells (Fig. 1C). Conversely, exogenous expression of EWS-FLI-1 in human mesenchymal stem cells, the putative cells of origin of Ewing sarcoma [1-5], induced pappalysin-1 transcript levels (Fig. 1D). Using chromatin immunoprecipitation, we detected the binding of EWS-FLI-1 to the papplysin-1 gene promoter in A673 and EW8 Ewing sarcoma cells, and this binding was abolished by shRNA-mediated silencing of EWS-FLI-1 (Fig. 2). These results indicate that papplysin-1 is a novel direct transcriptional target of EWS-FLI-1. Consistent with the activation of pappalysin-1 gene transcription by EWS-FLI-1, Ewing sarcoma tumors and cell lines expressed high levels of pappalysin-1 mRNA compared with mesenchymal stem cells (Fig. 3).

**Figure 1.**
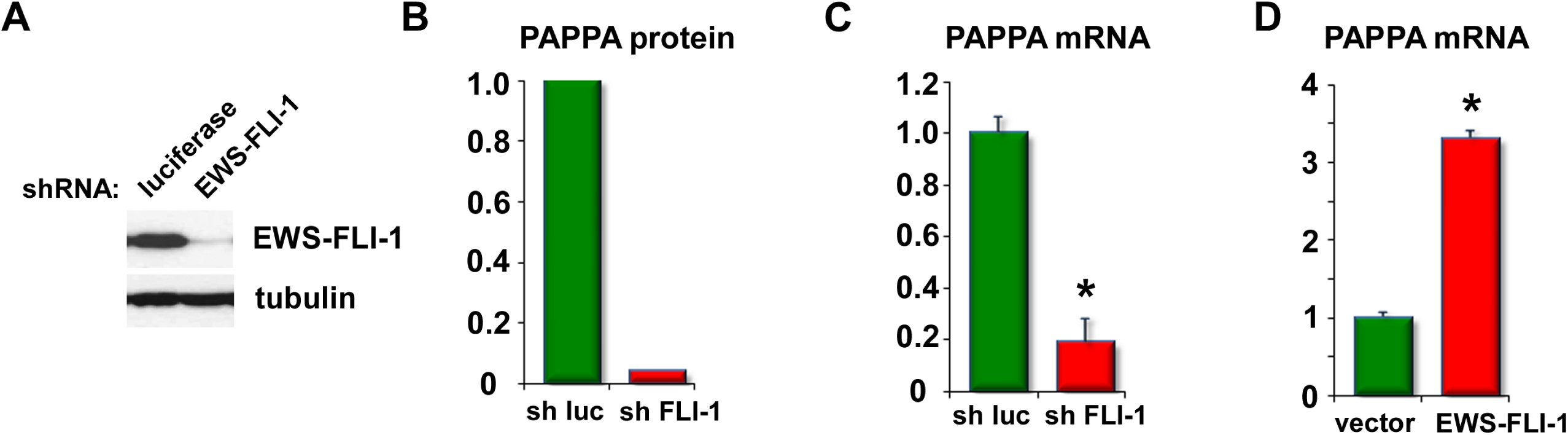
EWS-FLI-1 induces pappalysin-1 expression. (A) shRNA-mediated silencing of EWS-FLI-1 in A673 Ewing sarcoma cells. A673 cells were infected with lentiviruses expressing an shRNA against FLI-1 C-terminal region or luciferase (control) and were selected with 2 μg/ml puromycin for 2 days. Whole cell lysates were prepared and immunoblotting was performed using antibodies against FLI-1 C-terminus and tubulin (loading control). (B) Reduced pappalysin-1 protein levels in A673 cell secretome upon EWS-FLI-1 silencing. (C) EWS-FLI-1 silencing in A673 cells results in reduced pappalysin-1 transcript levels. EWS-FLI-1 was silenced as in (A) and the pappalysin-1 mRNA levels were examined by real-time PCR. An asterisk denotes p < 0.05 compared with luciferase shRNA-expressing cells. (D) EWS-FLI-1 expression in human mesenchymal stem cells results in increased pappalysin-1 transcript levels. Human mesenchymal stem cells were infected with lentiviruses expressing EWS-FLI-1 or empty vector and were selected with 2 μg/ml puromycin for 2 days. The pappalysin-1 mRNA levels were examined by real-time PCR. An asterisk denotes p < 0.05 compared with vector-expressing cells.

**Figure 2.**
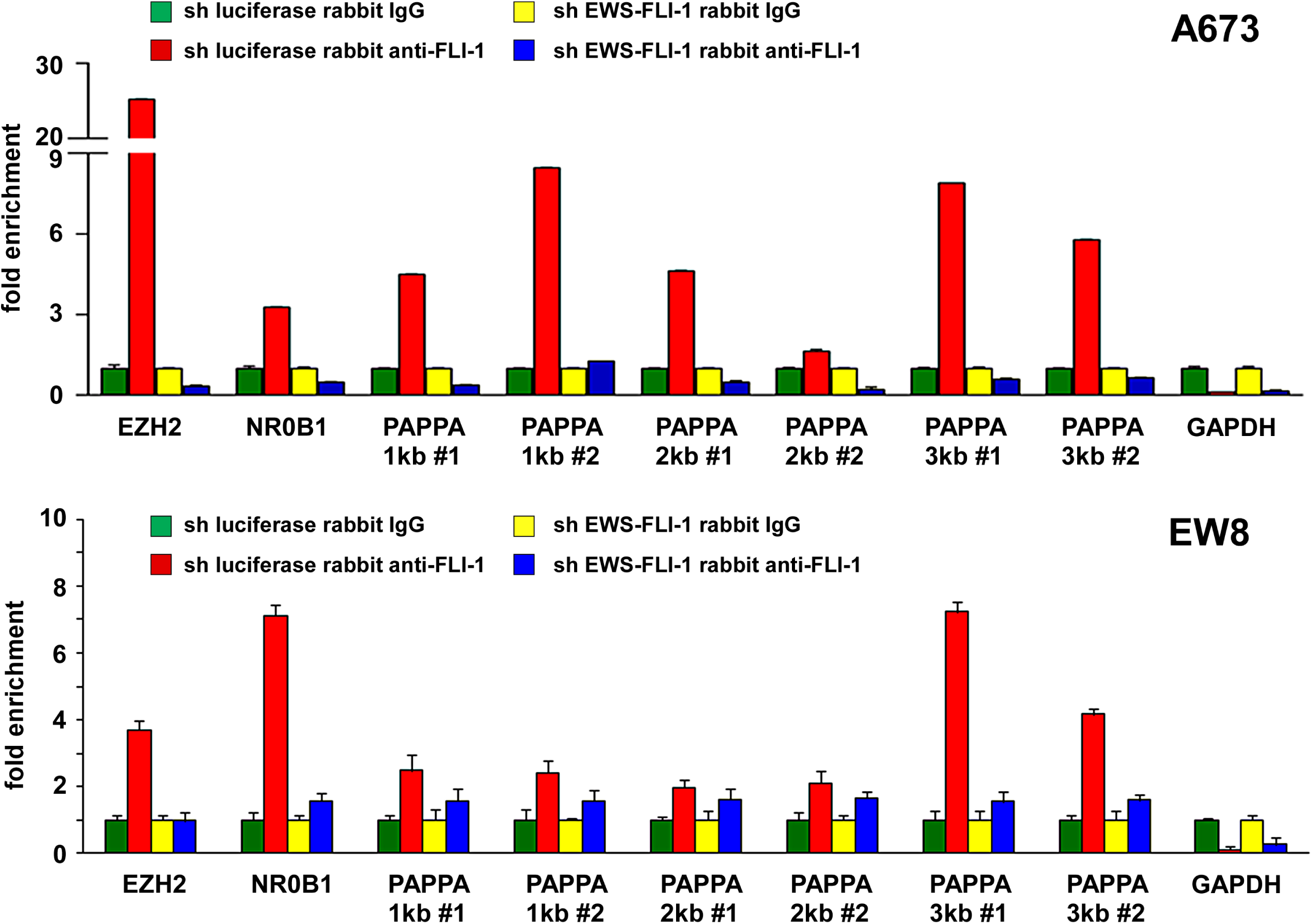
EWS-FLI-1 binds to the pappalysin-1 gene promoter in Ewing sarcoma cells. The binding of EWS-FLI-1 to the pappalysin-1 promoter was examined by chromatin immunoprecipitation in A673 and EW8 Ewing sarcoma cells. Two different primer pairs (#1 and #2) were used to amplify 1, 2, and 3 kb upstream regions. The specificity of the binding was verified by EWS-FLI-1 silencing. EZH2 [34] and NR0B1 [35] are known EWS-FLI-1 targets. GAPDH serves as a negative control.

**Figure 3.**
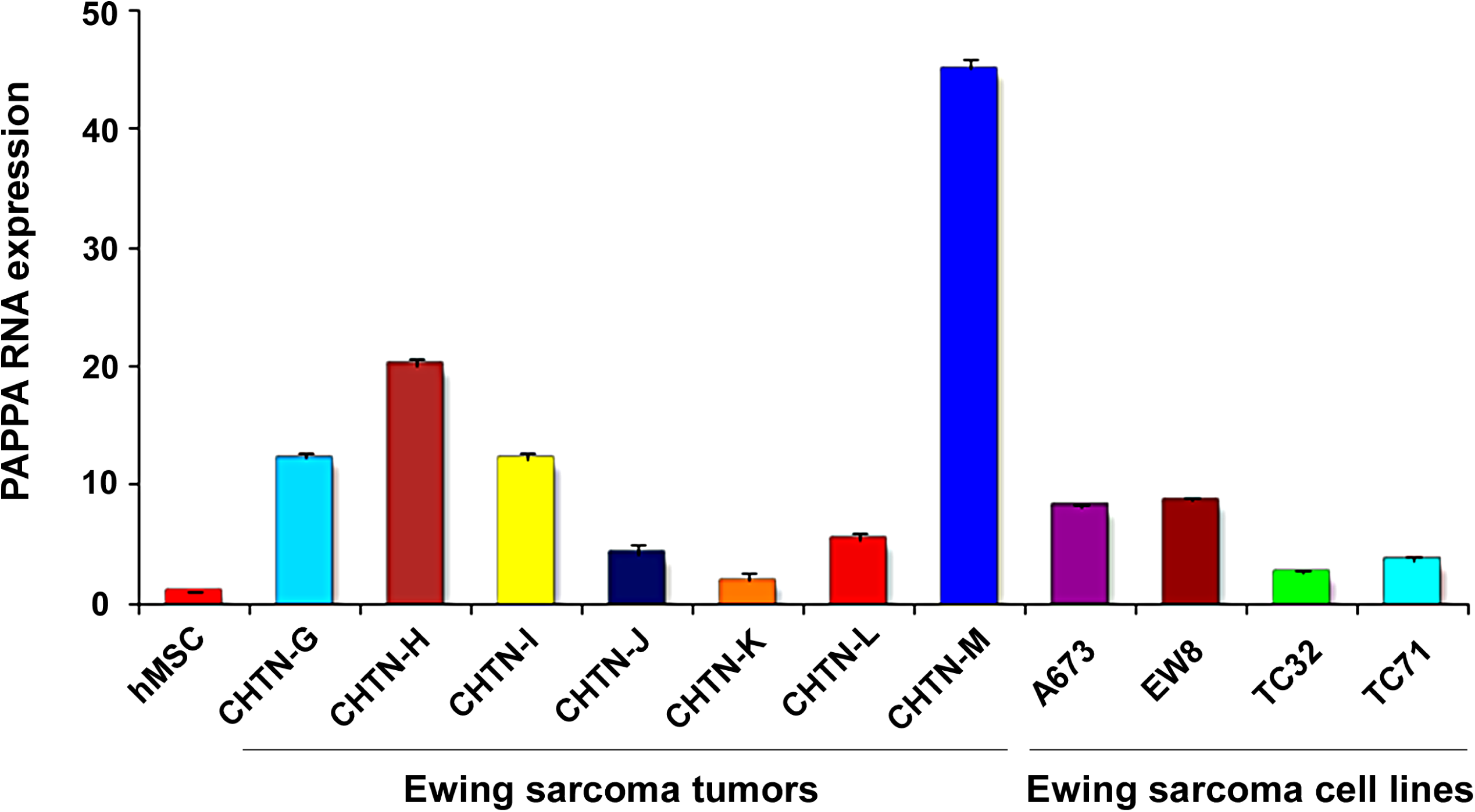
Pappalysin-1 is highly expressed in Ewing sarcoma tumors and cell lines. The pappalysin-1 mRNA expression in seven Ewing sarcoma tumor samples and four Ewing sarcoma cell lines was examined by real-time PCR. All RNA levels were normalized to the RNA levels in human mesenchymal stem cells.

### Silencing of pappalysin-1 results in accumulation of IGFBPs, reduced bioactive IGF-1, and reduced IGF signaling in Ewing sarcoma

Pappalysin-1 is a cell surface metalloproteinase that cleaves IGFBP2, IGFBP4, and IGFBP5 and releases IGF from inhibitory IGFBPs, leading to increased local bioavailability of IGF and enhanced IGF signaling [17, 18]. As shown in Fig. 4, silencing of pappalysin-1 by siRNA resulted in the accumulation of IGFBP2, IGFBP4, and IGFBP5 in the secretome of four Ewing sarcoma cell lines (A673, EW8, TC32, and TC71). Free bioactive IGF-1 levels dramatically declined in A673 and EW8 cell secretome upon papplysin-1 silencing (Fig. 5), which is consistent with the accumulation of IGFBPs (Fig. 4). Tyrosine auto-phosphorylation of IGF-1 receptor and serine 473 phosphorylation of Akt were significantly reduced upon pappalysin-1 silencing in all four Ewing sarcoma cell lines (Fig. 4). These results suggest that the proteolytic cleavage of IGFBPs by pappalysin-1 is necessary for the maintenance of IGF signaling in Ewing sarcoma.

**Figure 4.**
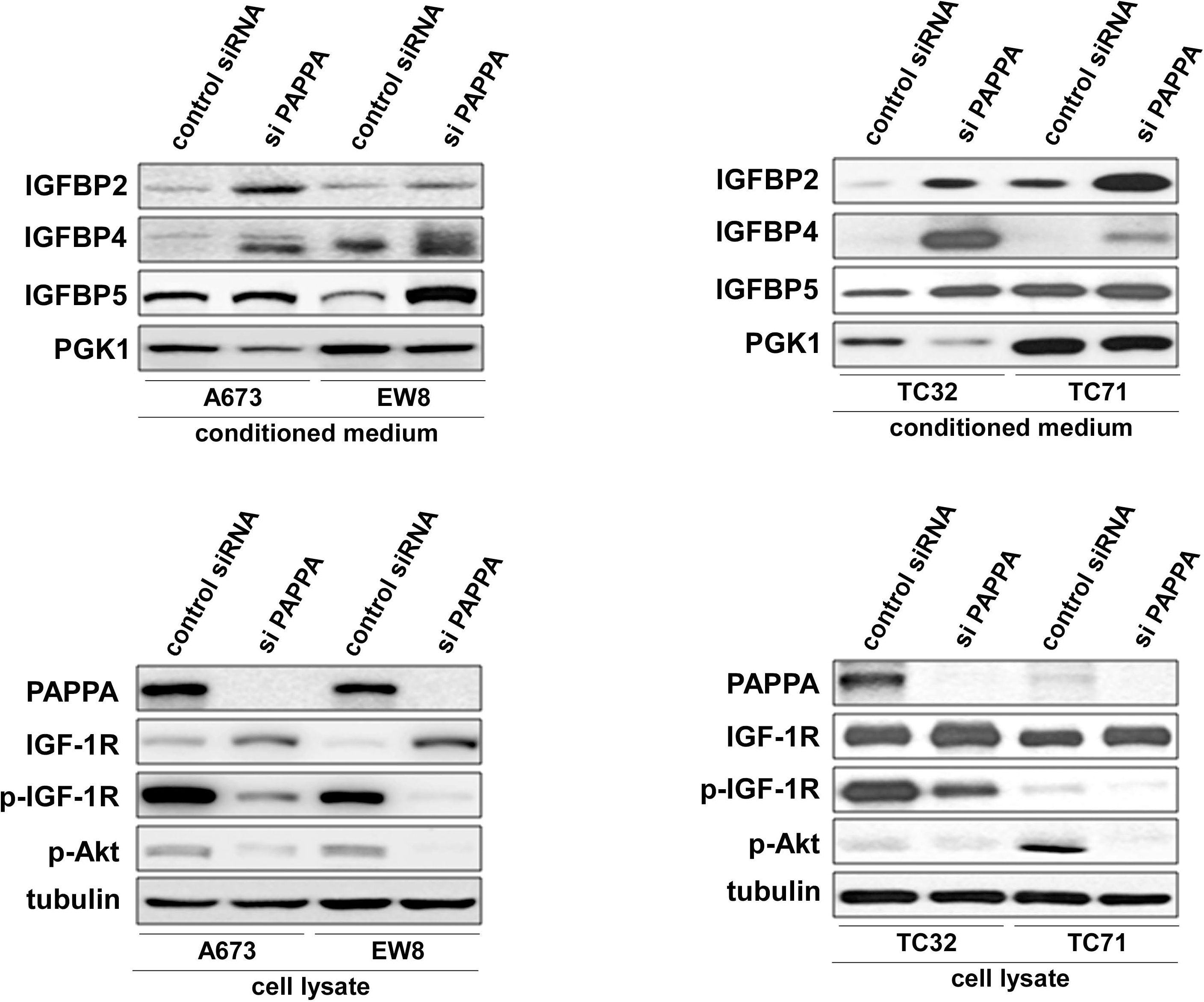
Pappalysin-1 silencing results in accumulation of IGFBPs and reduced IGF signaling in Ewing sarcoma cells. Pappalysin-1 expression was silenced by siRNA transfection in four Ewing sarcoma cell lines (A673, EW8, TC32, and TC71). Immunoblotting was performed using antibodies against IGFBP2, IGFBP4, IGFBP5, PGK1, pappalysin-1, IGF-1R, phospho-IGF-1R (phosphorylated at tyrosine 1135), phospho-Akt (phosphorylated at serine 473), and tubulin.

**Figure 5.**
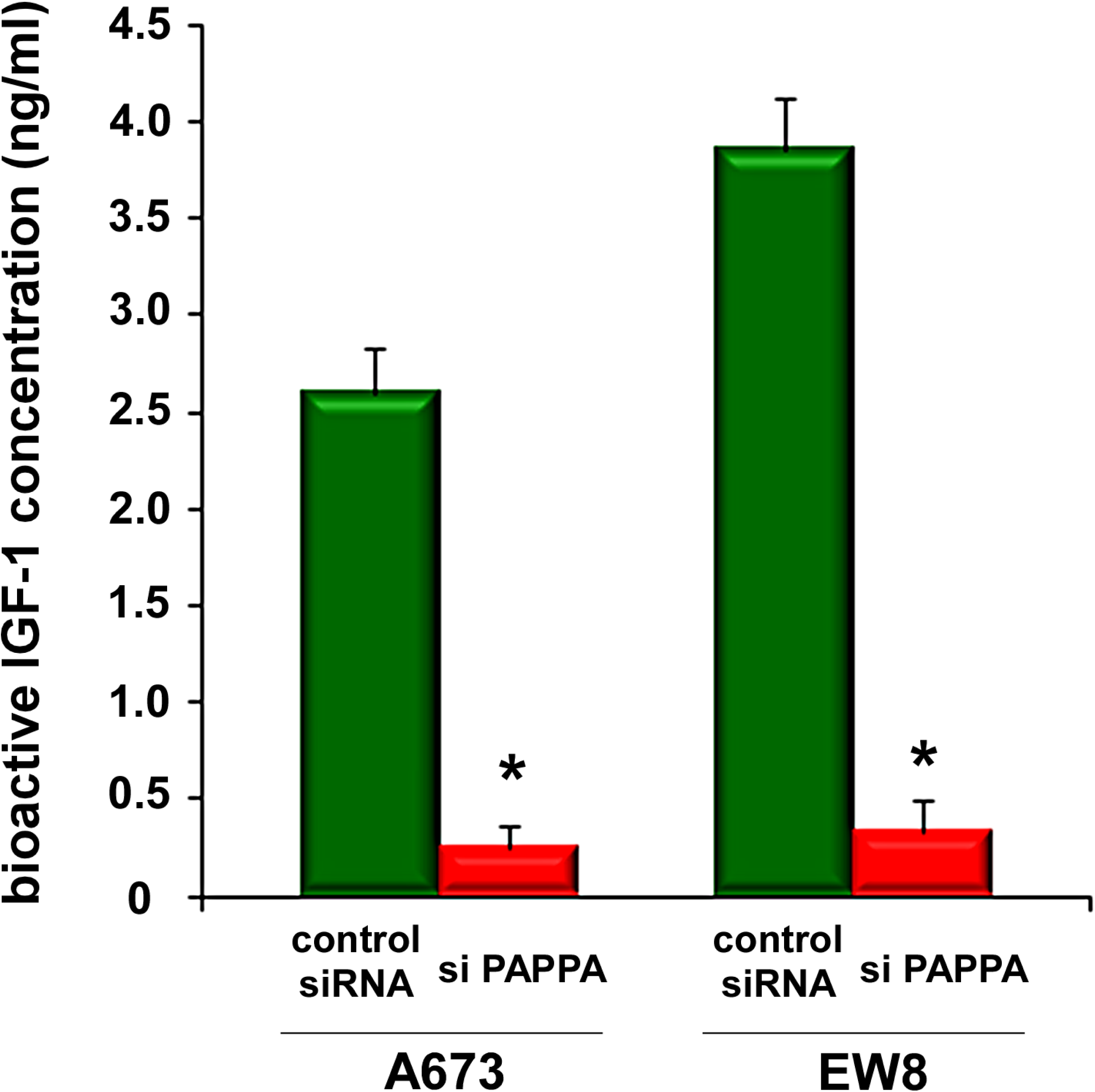
Pappalysin-1 silencing results in reduced bioactive IGF-1 levels in Ewing sarcoma secretome. Pappalysin-1 was silenced in A673 and EW8 cells as in Figure 4 and the levels of free bioactive IGF-1 were determined using bioactive IGF-1 ELISA. Asterisks denote p < 0.05 compared with control siRNA transfected cells.

### Silencing of pappalysin-1 inhibits anchorage-dependent and anchorage-independent growth and xenograft tumorigenicity of Ewing sarcoma cells

Pappalysin-1 silencing by siRNA or shRNA (Fig 6A) strongly inhibited the proliferation of Ewing sarcoma cells (Fig. 6B and 6D). Proliferation arrest induced by pappalysin-1 silencing was completely rescued by the addition of recombinant pappalysin-1 protein to the culture medium (Fig. 6C), indicating that Ewing sarcoma is dependent on extracellular pappalysin-1 protein.

**Figure 6.**
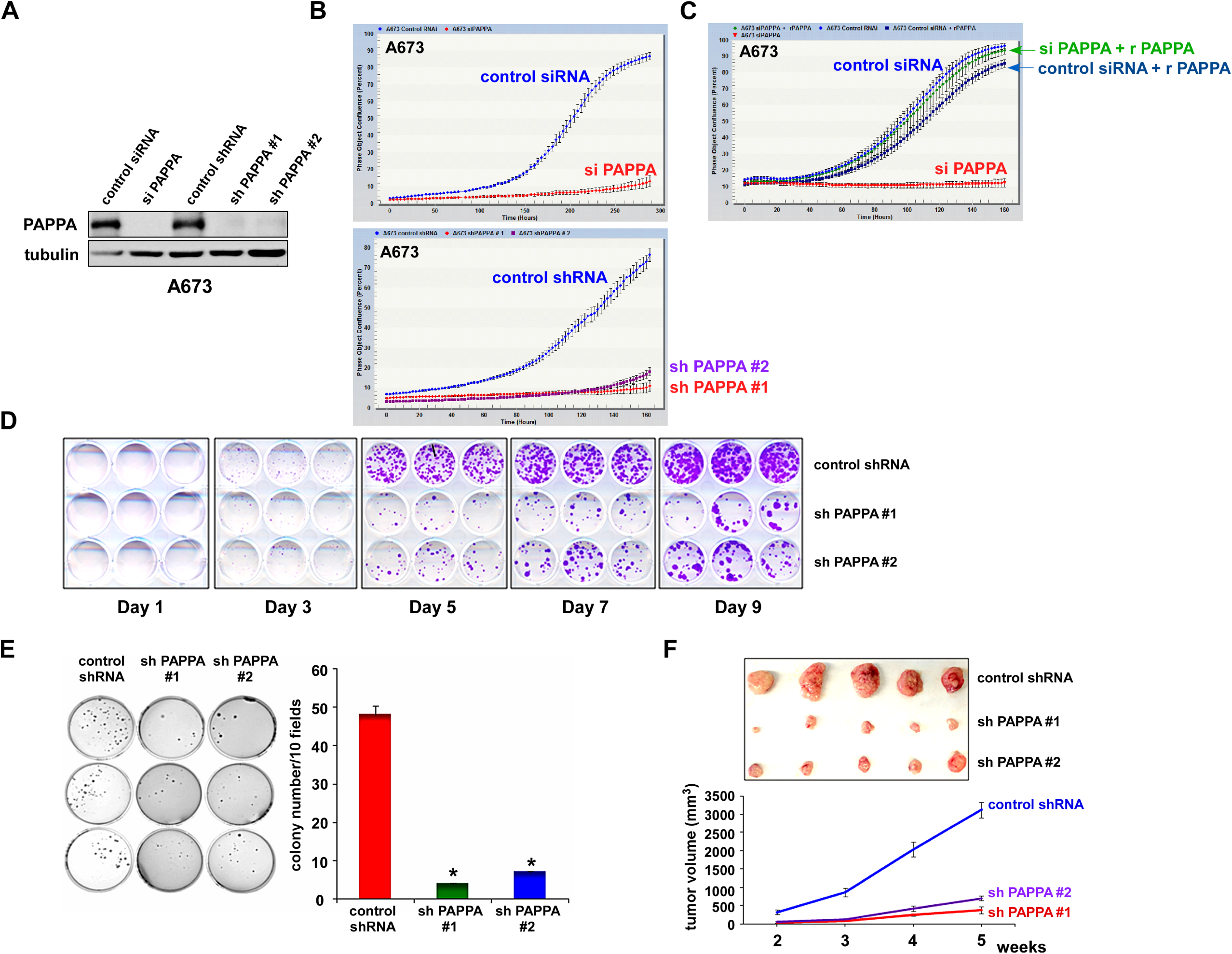
Pappalysin-1 silencing inhibits in vitro and in vivo growth of Ewing sarcoma cells. (A) Silencing of pappalysin-1 by siRNA and shRNA in A673 cells. (B) Pappalysin-1 silencing inhibits Ewing sarcoma cell proliferation. Pappalysin-1 was silenced by siRNA or shRNA and cell proliferation was assessed by using IncuCyte live-cell imaging system. (C) Recombinant pappalysin-1 protein rescues growth arrest induced by pappalysin-1 silencing.Sixteen hours after siRNA transfection, recombinant pappalysin-1 protein was added to the culture medium at 500 ng/ml where indicated and cell proliferation was assessed by IncuCyte system. (D) Crystal violet staining of A673 cells expressing pappalysin-1 shRNAs or control scrambled shRNA. (E) Pappalysin-1 silencing inhibits anchorage-independent growth of Ewing sarcoma cells.A673 were infected with lentiviruses expressing shRNAs against pappalysin-1 or control scrambled shRNA and were selected with 2 μg/ml puromycin. Four days after infection, cells were plated in semi-solid medium. Three week after culture, colonies were counted and photographed. Asterisks denote p < 0.05 compared with control shRNA-expressing cells.(E) Pappalysin-1 silencing inhibits xenograft tumorigenicity of Ewing sarcoma cells.A673 cells were infected with lentiviruses expressing pappalysin-1 shRNAs or control scrambled shRNA and were selected with 2 μg/ml puromycin for 2 days. Each cell type was subcutaneously injected into the flanks of SCID mice (2x10^6^ cells/injection, n=5). Tumor growth was monitored weekly using a caliper. The photograph of dissected tumors at 5 weeks after injection is shown on the top.

One of the hallmarks of cancer is the ability to proliferate independent of anchorage. Importantly, silencing of pappalysin-1 in A673 cells resulted in dramatic inhibition of soft agar colony formation (Fig. 6E), indicating that pappalysin-1 plays an essential role in anchorage-independent growth of Ewing sarcoma cells.

To test the role of pappalysin-1 in the tumorigenicity of Ewing sarcoma, we employed xenograft tumorigenicity assays in SCID mice. A673 cells were infected with lentiviruses expressing two different shRNAs against pappalysin-1 or control scrambled shRNA. After puromycin selection, cells were subcutaneously injected into the flanks of SCID mice. Tumor volume was determined using a caliper. As shown in Fig. 6F, pappalysin-1 silencing strongly inhibited xenograft tumor growth (p < 0.05).

Collectively, these results indicate that pappalysin-1 is required for *in vitro* and *in vivo* growth of Ewing sarcoma cells.

It is well established that IGF signaling plays an important role in Ewing sarcoma. High levels of IGF-1 and IGF-1R are expressed in Ewing sarcoma cell lines and tumors [10, 11] and inhibition of IGF-1R suppresses the xenograft tumorigenicity of Ewing sarcoma cells [12]. EWS-FLI-1 was shown to silence the expression of IGFBP3, an inhibitor of IGF signaling [16]. In this study, we uncovered that EWS-FLI-1 creates a cell surface microenvironment that is conducive to IGF signaling through direct transcriptional induction of pappalysin-1, a cell surface protease that cleaves IGFBP2, IGFBP4, and IGFBP5 (concept shown in Fig. 7). Our data suggest that pappalysin-1 stimulates IGF signaling in Ewing sarcoma by increasing the bioactive IGF levels in the vicinity of cell surface where IGF and IGF-1R interaction occurs. Downregulation of pappalysin-1 dramatically inhibited anchorage-dependent and anchorage-independent growth and xenograft tumorigenicity of Ewing sarcoma cells. These results indicate that despite high IGF-1 and IGF-1R expression and silencing of IGFBP3 by EWS-FLI-1, Ewing sarcoma is additionally dependent on pappalysin-1 to sustain IGF signaling and cell proliferation.

**Figure 7.**
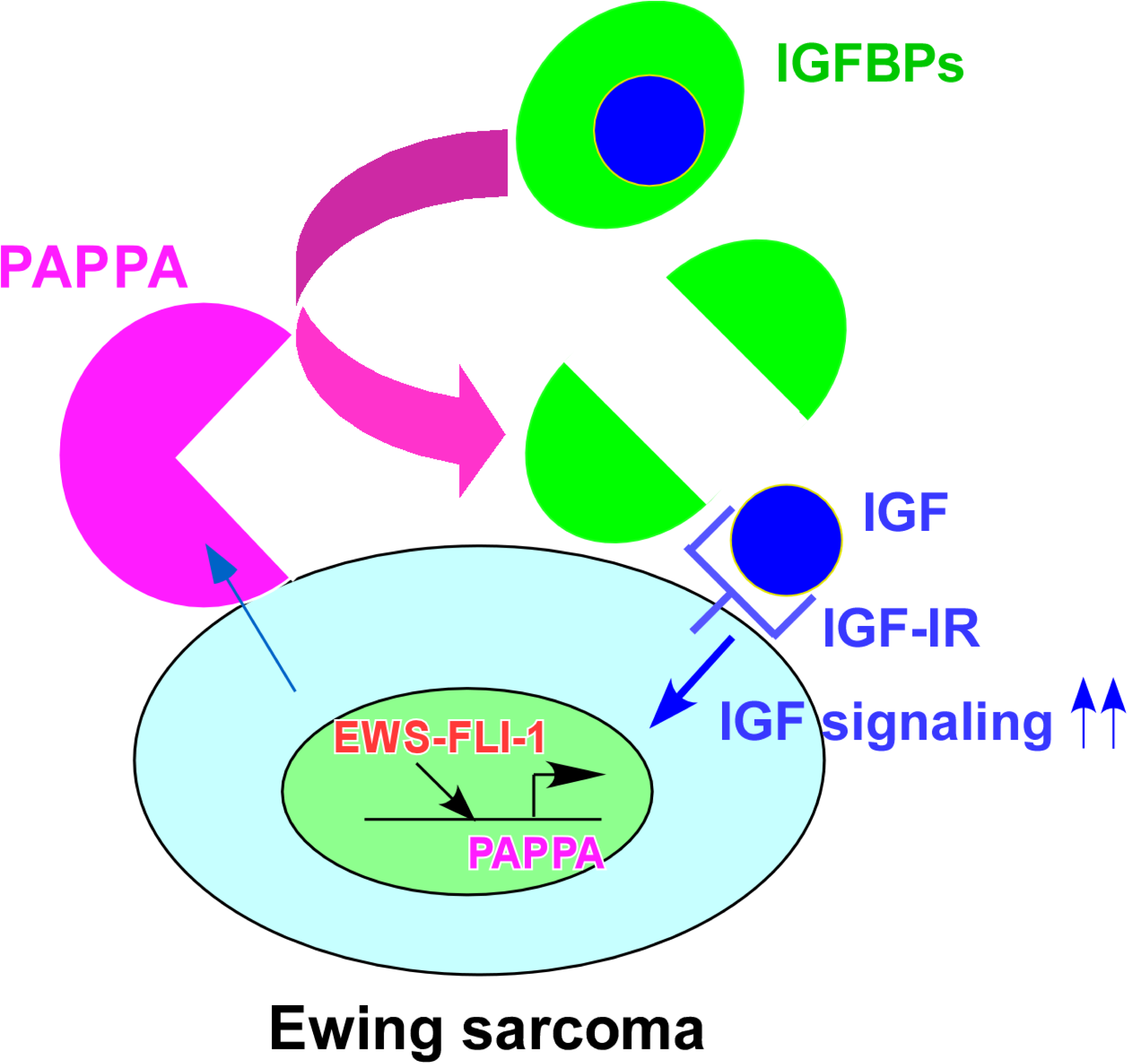
Model for the stimulation of IGF signaling by pappalysin-1 in Ewing sarcoma.EWS-FLI-1 directly activates the expression of pappalysin-1, which cleaves IGFBPs and increases bioactive IGF levels in the vicinity of cell surface, leading to enhanced IGF signaling and proliferation.

Important roles played by IGF signaling in Ewing sarcoma had generated high hopes for therapeutic targeting of this pathway, primarily by anti-IGF-1R antibodies. However, anti-IGF-1R clinical trials in Ewing sarcoma resulted in relatively low response rates [19-25]. The data presented in this paper suggest that downregulating pappalysin-1 to limit the cell surface bioavailability of IGF merits further investigation as an alternative to anti-IGF-1R antibodies. Additionally, a recent study identified pappalysin-1 as Ewing sarcoma tumor antigen and generated CD8^+^ T cells engineered to express TCR specific to pappalysin-1, which displayed anti-Ewing sarcoma effect in mouse xenograft models [26]. This study indicated that pappalysin-1 is a suitable antigen for cytotoxic T cell-mediated killing of Ewing sarcoma. Our data suggest that pappalysin-1 is a functionally important target in Ewing sarcoma, in addition to being a promising antigenic target for T cell-mediated immunotherapy. Therefore, it will now be important to determine the efficacy of pappalysin-1 targeting in Ewing sarcoma, both by immunotherapy and by pharmacological inhibition of pappalysin-1 activity.

## Materials & Methods

### Cell culture

A673 cells were cultured in Dulbecco’s modified Eagle’s medium (DMEM) supplemented with 10% fetal calf serum. EW8, TC71, and TC32 cells were cultured in RPMI 1640 medium supplemented with 10% fetal calf serum. 293T cells were cultured in DMEM supplemented with 10% calf serum. Cord blood-derived human mesenchymal stem cells were purchased from Vitro Biopharma (Golden, CO) and cultured in low serum MSC-GRO following the manufacturer’s procedure. Calcium phosphate co-precipitation was used for transfection of 293T cells. Lentiviruses were prepared by transfection in 293T cells following System Biosciences’ protocol and the cells infected with lentiviruses were selected with 2 μg/ml puromycin for 48 hours as described [27, 28]. The target sequences for shRNAs are as follows: FLI-1 C-terminus shRNA, AACGATCAGTAAGAATACAGAGC; luciferase shRNA, GCACTCTGATTGACAAATACGATTT; pappalysin-1 shRNA-1, CGGACAGACATTGTGTGACAA; pappalysin-1 shRNA-2, GCTGTATGACAAATGTTCTTA; and scrambled shRNA, CCTAAGGTTAAGTCGCCCTCG. The following siRNAs were used: human pappalysin-1 siRNA SMARTpool (M-005130-02-0005, Dharmacon) and Non-Targeting siRNA Pool #2 (D-001206-14-05, Dharmacon). siRNA transfection was performed using Lipofectamine™ RNAiMAX Transfection Reagent (Thermo Fisher). Recombinant pappalysin-1 protein (2487-ZNF-020) was purchased from R&D Systems.

### Protein sample preparation and proteomic analysis

The preparation of secreted protein samples, GeLC-MS/MS analysis, and proteomics data analysis were performed essentially as described [29]. A673 cells were infected with lentiviruses expressing an shRNA against FLI-1 C-terminal region or luciferase (control) and were selected with 2 μg/ml puromycin for 2 days. Cells were washed six times with DMEM without serum. Subsequently, cells were cultured in DMEM without serum for 24 hours and the culture supernatant was harvested. The supernatant was centrifuged, filtered through a 0.45 μm filter (Millipore), and concentrated using a 3,000 Dalton cut-off Amicon Ultra Centrifugal Filter Units (Millipore). The proteins in each sample were fractionated by SDS-PAGE and visualized by Coomassie blue. Each gel lane was divided into six slices, and the proteins in each slice were digested in situ with trypsin (Promega modified) in 40 mM NH4HCO3 overnight at 37 °C. The resulting tryptic peptides were analyzed by HPLC-ESI-tandem mass spectrometry (HPLC-ESI-MS/MS) on a Thermo Fisher LTQ Orbitrap Velos mass spectrometer. The Xcalibur raw files were converted to mzXML format using ReAdW and were searched against the UniProtKB/Swiss-Prot human protein database using X! Tandem. The X! Tandem search results were analyzed by the Trans-Proteomic Pipeline [30] version 4.3. Peptide/protein identifications were validated by Peptide/ProteinProphet [31, 32].

### RNA samples and quantitative Real-Time PCR

De-identified Ewing sarcoma tumor RNA samples were obtained from the Cooperative Human Tissue Network. Total cellular RNA was isolated using TRIzol reagent (Invitrogen). Reverse transcription was performed using High Capacity cDNA Reverse Transcription Kit (Thermo Fisher) as per manufacturer’s instructions. Quantitative PCR was performed using PowerUp SYBR Green Master Mix (Thermo Fisher) on Applied Biosystems ViiA 7 Real-Time PCR System. Each sample was analyzed in triplicate. The following primers were used: pappalysin-1 5’ primer, CAGAATGCACTGTTACCTGGA, 3’ primer, GCTGATCCCAATTCTCTTTCA; GAPDH 5’ primer, GGTGTGAACCATGAGAAGTATGA, 3’ primer, GAGTCCTTCCACGATACCAAAG.

### Immunoblotting, ELISA, and antibodies

Immunoblotting was performed as described [27, 28]. The levels of free bioactive IGF-1 were determined using IGF-I Bioactive ELISA (AnshLabs). The following antibodies were used: rabbit polyclonal anti-phospho-Akt (9271, Cell Signaling Technologies); rabbit polyclonal anti-FLI-1 (ab15289, Abcam); rabbit polyclonal anti-IGFBP2 (3922, Cell Signaling Technologies); goat polyclonal anti-IGFBP4 (AF804, R&D Systems); mouse monoclonal anti-IGFBP5 (sc-515116, Santa Cruz Biotechnology); rabbit monoclonal anti-IGF-1R (9750, Cell Signaling Technologies); rabbit monoclonal anti-phospho-IGF-1R (3918, Cell Signaling Technologies); goat polyclonal anti-PGK1 (sc-17943, Santa Cruz Biotechnologies); and mouse monoclonal antitubulin (DM1A, Thermo Scientific);

### Chromatin immunoprecipitation

Chromatin immunoprecipitation (ChIP) was performed as described [33] using rabbit polyclonal anti-FLI-1 antibody (ab15289, Abcam) or control rabbit IgG (ab37415, Abcam). The primer sequences used for ChIP are as follows: pappalysin-1 1kb #1 5’ primer, GGAGGAGTTGGGCTGTATTT, 3’ primer, CTTCGCTTCTTCACCCTTCT; pappalysin-1 1 kb #2 5’ primer, TTAGCTGAAGCCAGCCTTATC, 3’ primer, CCCTTTACCTCTTTCCCTCTTC; pappalysin-1 2 kb #1 5’ primer, GGAGCAGCTCGGAAGATAAG, 3’ primer, TGTGTGAAGTTTGCATGTGAAT; pappalysin1 2kb #2 5’ primer, GCCTAATTGCCACAACTGAAG, 3’ primer, TGCAACTCCAAGTCATCTGTAA; pappalysin-1 3kb #1 5’ primer, CCAGAGCATATCTTGTCCTCAAA, 3’ primer, CATTTCCTACTCCCTCCAACAC; pappalysin-1 3kb #2 5’ primer, CACAAAGCAGAATAAGATCCTGAG, 3’ primer, TGTATCATGTACTGCTATCCCTTT; EZH2 5’ primer, GACACGTGCTTAGAACTACGAACAG, 3’ primer TTTGGCTGGCCGAGCTT; NR0B1 5’ primer, GTTTGTGCCTTCATGGGAAATGGTTATTC, 3’ primer, CTAGTGTCTTGTGTGTCCCTAGGG; GAPDH 5’ primer, TCCTCCTGTTTCATCCAAGC, 3’ primer, TAGTAGCCGGGCCCTACTTT.

### Cell proliferation assays

Anchorage-dependent cell proliferation was assessed by IncuCyte live-cell imaging system and by crystal violet staining.

Anchorage-independent cell proliferation was evaluated by soft agar colony formation assays. A673 cells were infected with lentiviruses expressing shRNAs against pappalysin-1 or scrambled shRNA and were selected with 2 μg/ml puromycin. Four days after infection, 4x10^3^ cells were plated in soft agar. The soft agar cultures were comprised of two layers: a base layer (4 ml in a 60 mm dish; DMEM/10% fetal calf serum/0.6% noble agar (A5431, Sigma-Aldrich)/ penicillin/streptomycin) and a cell layer (2 ml in a 60 mm dish; DMEM/10% fetal calf serum/0.3% noble agar/penicillin/streptomycin). Colonies were grown for three weeks and counted. Colonies (>50 cells) were scored by randomly counting 10 fields per dish.

### Xenograft tumorigenicity assays

A673 cells were infected with lentiviruses expressing pappalysin-1 shRNAs or scrambled shRNA and were selected with 2 μg/ml puromycin for 2 days. Each cell type was subcutaneously injected into the flanks of SCID mice (2x10^6^ cells/injection, n=5). Tumor growth was monitored weekly using a caliper.

### Statistical analysis

Statistical significance was calculated by using Graphpad Prism software (version 6.0f) with a two-tailed Student’s t test.

## Acknowledgements

We thank the Cooperative Human Tissue Network for Ewing sarcoma tumor RNA samples and the UTHSCSA Institutional Mass Spectrometry Laboratory for mass spectrometry analysis. This work was supported by the Owens Medical Research Foundation (to Y.S.), by the National Cancer Institute, the National Institutes of Health (CA202485 to Y.S; CA165995 to P.J.H.), by the Cancer Prevention and Research Institute of Texas (RP160487 and RP160841 to Y.S.; RP160716 to P.J.H.), by the National Center for Advancing Translational Sciences, National Institutes of Health, through the Clinical and Translational Science Award (CTSA) UL1 TR001120, and by the CTRC P30 Cancer Center Support Grant from the National Cancer Institute (CA054174).

